# Cyclo olefin polymer-based solvent-free mass-productive microphysiological systems

**DOI:** 10.1101/2020.06.24.170035

**Authors:** Makoto Yamanaka, Xiaopeng Wen, Satoshi Imamura, Risako Sakai, Shiho Terada, Ken-ichiro Kamei

## Abstract

A microphysiological system (MPS) holds a great promise for drug screening and toxicological testing as an alternative to animal models. However, this platform has several issues in terms of the materials used (e.g., polydimethylsiloxane), such as the absorbance of tested drug candidates and fluorescent dyes by the material, as well as the effect on cultured cellular status, thus misleading the results obtained from cell assays and fabrication processes. Hence, to eliminate the issues mentioned above, we developed a cyclo olefin polymer (COP)-based MPS via photobonding process using vacuum ultraviolet (VUV), named COP-VUV-MPS. COP-VUV-MPS showed better chemical resistance and avoided molecule absorption. COP-VUV-MPS could maintain the stemness of environmentally sensitive human-induced pluripotent stem cells without causing undesired cellular phenotypes and gene expression. These results suggested that COP-VUV-MPS might be broadly used for the advancement of MPS and applications in drug development and in vitro toxicological testing.

## Introduction

Microphysiological systems (MPSs)^1–3^ or Organ/Body-on-a-Chip (O/BoC) systems^4–7^ hold great potential for application in drug development and toxicological testing. These platforms recapitulate physiological and pathological conditions *in vivo* in a chip without the use of actual organs, as an alternative to animal testing.^2,3^

Most MPS are based on microfluidic technology, since it offers several benefits over the conventional macroscopic cell culture settings, such as precise liquid control, designed extracellular geometry, cellular positioning, and culturing in two- and three-dimensional fashions, which are very important for mimicking *in vivo* organ structures.^8,9^ However, there is a critical issue to be solved. It is well-known that polydimethylsiloxane (PDMS), which has been widely used for a long time for fabricating microfluidic devices for MPS platforms due to its biocompatibility, gas permeability, and transparency, has several limitations. These include hydrophobicity and porosity, leading to chemical/drug absorption, water evaporation, deformation, and leaching un-crosslinked reagents.^10–12^ These limitations might cause some concerns for cellular experiments, e.g., reduction of tested drugs/chemicals, evaporation of cell culture media and unexpected cellular behaviors. The use of PDMS is also challenging for mass production and long-term storage of the fabricated MPS. Therefore, there is a clear need to find alternatives of PDMS, but we have not come across suitable materials to solve these issues for over a decade.

While seeking for alternatives to PDMS, we have realized that cyclo olefin polymer (COP) is advantageous for MPSs, not only over PDMS but also glass, polystyrene, and polymethyl methacrylate.^13–15^ COP is an amorphous polymer and shows chemical/physical stability, high purity and optical clarity, and has been used for a variety of applications, including not only displays and packaging films but also medical products. Since COP also shows thermostability with high modulus, metal molding process can be used for mass production of MPSs without deformation of microstructures and should be beneficial for MPS applications. However, although COP shows high purity, the fabrication process of a COP microfluidic device often requires solvents and tapes which might affect cellular phenotypes, dismissing the advantages of COP.^16^ Therefore, an alternative way of fabricating a COP microfluidic device needs to be developed.

Here, we developed a COP-based MPS without the use of PDMS and solvents for the fabrication process (**Fig. 1a**). Our proposed MPS provides (1) stable microstructures, (2) chemical resistance to organic solvents and acids, (3) reduction of chemical compounds absorption, (4) no leakage and (5) applicable for mass production (**Fig. 1b**). To evaluate the effects on the cells cultured in the device, we used human pluripotent stem cells (hPSCs; i.e., embryonic stem cells [hESCs]^17^ and induced pluripotent stem cells [hiPSCs]^18,19^), which serve as a unique and powerful model, since they are susceptible to the external environment and stimuli.^20^

**Fig. 1.**
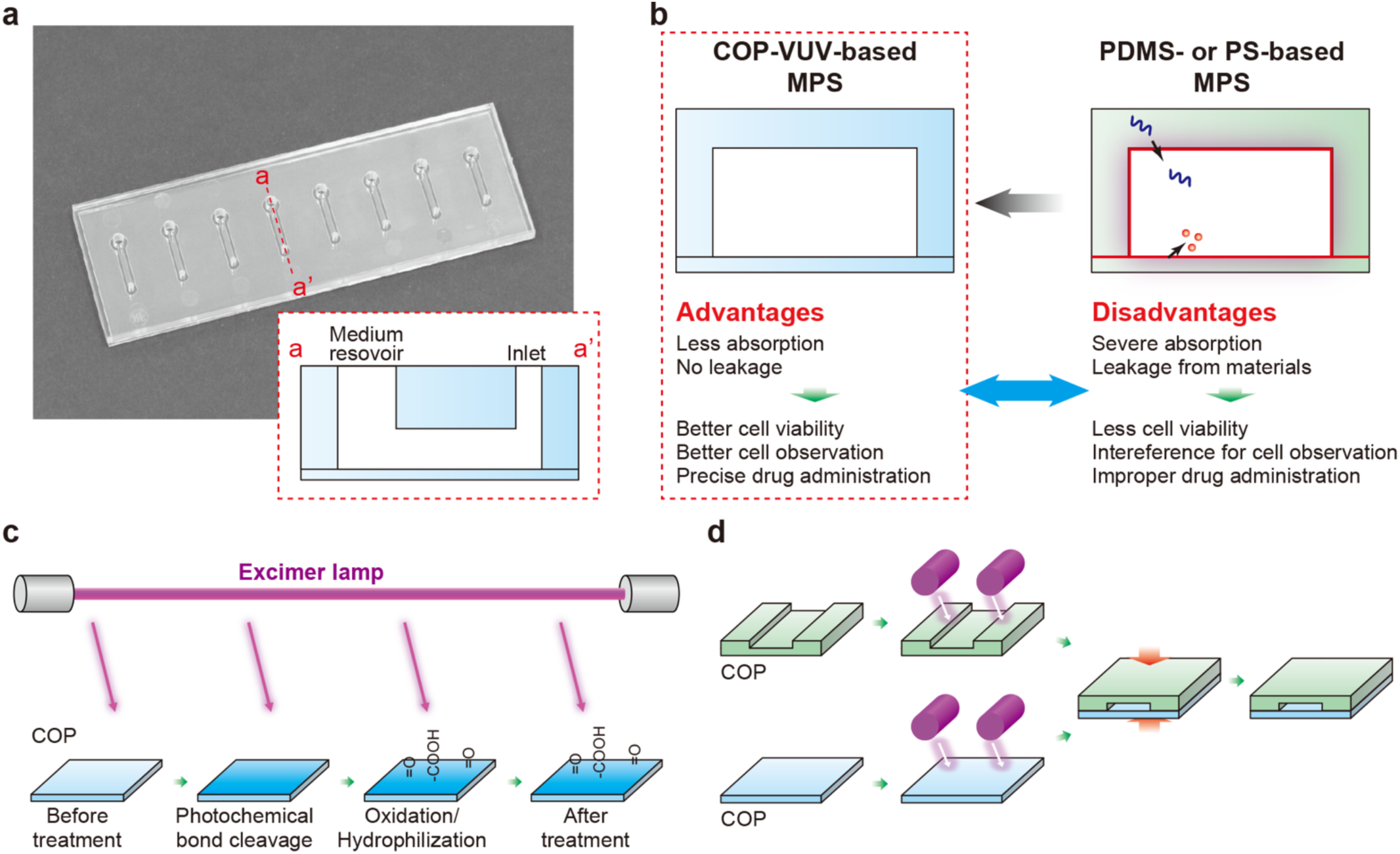
Overview of a polydimethylsiloxane (PDMS)- and solvent-free microphysiological system (MPS). (a) Photographs of actual cyclo olefin polymer (COP)-based MPS, and the illustration of one of the microfluidic channels. The device consists of two layers of COP and has eight microfluidic cell-culture channels (800 μm × 7500 μm × 250 μm; total volume in each channel is 13.2 μL) with a medium inlet and outlet (2 and 1 mm in diameters, respectively). (b) Advantages of a COP-vacuum ultraviolet (VUV) device over the conventional MPS comprising materials such as PDMS and polystyrene (PS). (c) VUV photobonding allows us to eliminate the limitations of MPSs. VUV light generated from an excimer lamp has a 172-nm wavelength, which cleaves the COP surface, resulting in oxidation and hydrophilization of the surface. (d) Schematic illustration of the fabrication of a COP microfluidic device with metal molding followed by VUV photobonding.

## Experimental

### Fabrication of COP-VUV-MPS

To accomplish aforementioned requirements for advancement of MPS, we used the vacuum ultraviolet (VUV) photobonding process for fabricating a COP-based MPS (namely COP-VUV-MPS, **Fig. 1c,d**).^21^ The VUV photobonding from an excimer light at a 172-nm wavelength was used, and this treatment generated the functional groups (e.g., hydroxy and carboxyl groups) for assembling two COP materials with heat treatment. COP-VUV-MPS comprises two layers of COP and eight microfluidic cell culture channels (800 μm in width, 7500 μm in length, 250 μm in height) with a medium inlet and outlet (2 and 1 mm in diameter, respectively), and the total volume in each channel is 13.2 μL. Two metal molds were used to fabricate the microfluidic structures. COP material (Zeonex 690R, Zeon, Tokyo, JAPAN) was injected to the two molds individually. After taking the structured COP components from the molds, the components were irradiated with VUV from an excimer lamp (172 nm; Ushio INC., Tokyo, JAPAN) at 25°C. The component surfaces were assembled using a heat press at less than 132°C. Finally, ethylene oxide gas (Japan Gas Co. Ltd., Kanagawa, JAPAN) was used for the decontamination of the fabricated COP-VUV-MPS.

### Fabrication of PDMS device

Polymethyl methacrylate (PMMA, Sumitomo Chemical Co., Ltd., Tokyo, JAPAN) was used following the micromilling procedure (CNC JIGBORER YBM 950V, Yasda Precision Tools K.K, Okayama, JAPAN) to fabricate the mold for a PDMS-MPS. Pre-PDMS (Sylgard184; A and B) was mixed at a 10:1 ratio and degassed for 1 h at 25°C. The mixed pre-PDMS was poured on the PMMA mold and then placed in an oven at 65°C for 2 h. The solidified PDMS was peeled off from the mold, and placed in an oven at 80°C for 24 h. The surface of PDMS was soaked in ethanol (EtOH) and sonicated to clean. PDMS and a glass slide (Matsunami Glass, Osaka, JAPAN) were treated with VUV from an excimer lamp (172 nm, Ushio INC.) at 25°C and then assembled at 80°C for 2 days. Before cell culture, the obtained PDMS device was irradiated with UV for decontamination.

### Evaluation of device deformation by solvents

To evaluate the deformability of COP-VUV-MPS, acids [1 M HCl aqueous solution (FUJIFILM Wako Pure Chemical Corp., Osaka, JAPAN)], bases [1 M NaOH solution (FUJIFILM Wako Pure Chemical Corp.)] and organic solvents [dimethyl sulfoxide (DMSO) 99.0% (w/w) (FUJIFILM Wako Pure Chemical Corp.), isopropyl alcohol (IPA) 99.5% (w/w) (NACALAI TESQUE, INC., Kyoto, JAPAN), EtOH 99.5% (w/w) (FUJIFILM Wako Pure Chemical Corp.) and acetonitrile 99.5% (w/w) (FUJIFILM Wako Pure Chemical Corp.)] were used to treat the device. Fluorescence microscopy was used to measure deformation of the microfluidic channels. Before the deformation test, 1 μg mL^-1^ fluorescein solution (Sigma Aldrich, St. Louis, MO, USA) was introduced into the microfluidic channel to measure the original width of the channel. Then, the channel was observed with a fluorescence microscope (BZ-X800, Keyence, Osaka, JAPAN) and measured using image analysis (ImageJ, ver.1.52p). The flow path was then completely replaced with the test solvent and treated at 20°C for 1 h. The flow path was measured again to determine the width of the channel.

### Evaluation of molecule absorption

The fluorescent imaging system (Typhoon FLA 7000, GE Healthcare, Chicago, IL; PMT sensitivity 500; Range mode L5) was used to measure the absorption of the tested substrates. AdipoRed (PT-7009, Lonza, Basel, Switzerland) and doxorubicin hydrochloride (DXR; Sigma Aldrich) were used as typical hydrophobic fluorescent molecules. AdipoRed original solution was diluted in phosphate-buffered saline (PBS) 40 times as a working solution. The AdipoRed working solution was incubated with the tested substrates at 37°C for 30 min and then washed with air blow, followed by fluorescent imaging. DXR was dissolved in DMSO to make 20-mM stock solution, and further dilution with PBS was performed to make a 20-μM working solution. The DXR working solution was incubated with the tested substrates at 37°C for 24 h, and then washed with air blow, followed by fluorescent imaging. We measured the fluorescence signals of the AdipoRed hydrophobic lipid fluorescent indicator [ex. 553 nm; em. 637 nm] and DXR anti-cancer drug [ex. 470 nm; em. 585 nm]) in the tested materials, using GE Typhoon FLA7000 fluorescent scanner.

### hiPSC culture

253G1 hiPSCs were obtained from RIKEN Bioresource Center. Before culturing, hESC-certified Matrigel (BD Bioscience, CA, USA) was diluted with Dulbecco’s modified Eagle medium (DMEM)/F12 medium (Sigma Aldrich) to 1.3% (v/v) and used to coat a culture dish. Matrigel was incubated in a dish for 24 h at 4°C. Then, excess Matrigel was removed, and the coated dish was washed with fresh DMEM/F12 medium.

TeSR-E8 medium (Stem Cell Technologies, Vancouver, Canada)^22^ was used for daily culturing of hiPSCs. For passaging, cells were dissociated with TrypLE Express (Life Technologies, Carlsbad, CA, USA) for 3 min at 37°C and harvested. A cell strainer was used to remove undesired cell aggregates from the cell suspension, and cells were centrifuged at 200 × *g* for 3 min and resuspended in TeSR-E8 medium. Cells were counted using a NucleoCounter NC-200 (Chemetec, Baton Rouge, LA, USA). TeSR-E8 medium containing 10 μM ROCK inhibitor Y-27632 (FUJIFILM Wako Pure Chemical Corp.) was used to prevent apoptosis of dissociated hPSCs on day 1.^23^ TeSR-E8 medium without the ROCK inhibitor was used on subsequent days, with daily medium changes.

### hiPSC culture in MPS

hiPSCs were cultured in the MPS as previously described.^24^ Briefly, before use, the MPS was wiped with 70% (v/v) EtOH and placed under UV light in a biosafety cabinet for over 30 min. Then, 10 μL of 1.3% (v/v) Matrigel in DMEM/F12 was introduced into each microfluidic cell culture channel and kept at 4°C for over 24 h. Then, excess Matrigel was removed, and coated channels were washed with fresh DMEM/F12 and TeSR-E8 medium.

hiPSCs cultured in a dish were harvested with 1 mL of TrypLE Express for 5 min at 37°C, then transferred into a 15-mL centrifuge tube. Four milliliters of TeSR-E8 medium was added into the tube, and hPSCs were centrifuged at 200 × *g* for 3 min. After decanting the supernatant, hPSCs were resuspended in 5 mL of TeSR-E8 medium and centrifuged at 200 × *g* for 3 min. Then, hPSCs were resuspended in TeSR-E8 medium supplemented with 10 μM Y-27632 and introduced into the MPS microfluidic channel at 2.0 × 10^6^ cells in 10 μL. Two hours after cell seeding, fresh TeSR-E8 supplemented with 10 μM Y-27632 was introduced into the microfluidic cell culture channel to remove debris. Then, the medium was changed every 12 h until further experiments.

### RNA purification

Cells cultured in the MPS channel were washed with 15 μL of D-PBS twice. Then, 10 μL of the RLT buffer in the RNeasy micro kit (Qiagen, Hilden, Germany) was introduced into the channel to make the cell lysate. The lysate was obtained from the channel and transferred into a 1.5-mL tube. Total RNA was isolated using RNeasy micro, following the manufacturer’s protocol. Purified RNA was quantified using Bioanalyzer 2100 (Agilent Technologies, Santa Clara, CA) with the Agilent RNA6000 pico chip.

### Quantitative RT-PCR

Total RNA was reverse transcribed to generate cDNA using PrimeScript RT master mix. A reaction mixture (21 μL) containing 20 ng cDNA, 12.5 μL SYBR Premix Ex Taq II (Tli RNaseH Plus; TaKaRa Bio, Shiga JAPAN), and 0.5 μL ROX reference dye was introduced into the wells of MicroAmp® Optical 96-Well Reaction Plate (Life Technologies), according to the manufacturer’s instructions. PCR conditions included an initial incubation at 95°C for 30 s, followed by 40 cycles of 95°C for 5 s, and then 60°C for 31 s on an Applied Biosystems 7300 real-time PCR system.

### Immunocytochemistry

Cells were fixed with 4% (v/v) paraformaldehyde in PBS for 20 min at 25°C and then permeabilized with 0.5% (v/v) Triton X-100 in PBS for 16 h at 25°C. Subsequently, cells were blocked in PBS (5% (v/v) normal goat serum, 5% (v/v) normal donkey serum, 3% (v/v) bovine serum albumin, 0.1% (v/v) Tween-20) at 4°C for 16 h and then incubated at 4°C for 16 h with the primary antibodies ([anti-human OCT3/4 mouse Ig, 1:100; Santa Cruz Biotechnology, Inc., USA]) in PBS with 0.5% (v/v) Triton X-100. Cells were then incubated at 37°C for 60 min with a secondary antibody (AlexaFluor 488 Donkey anti-rabbit IgG, 1:1000; Jackson ImmunoResearch, West Grove, PA, USA) in blocking buffer prior to a final incubation with 300 nM of 4‖,6-diamidino-2-phenylindole (DAPI) at 25°C for 30 min.

### Apoptotic cell staining

Annexin V staining was performed according to the product manual of Alexa Fluor 594-Annexin V conjugate (Molecular Probes, Eugene, OR). Briefly, after washing with Annexin-binding buffer (10 mM HEPES (pH 7.4), 140 mM NaCl, 2.5 mM CaCl_2_), the cells were stained with Alexa Fluor 594-Annexin V conjugate at 25°C for 15 min. Following cell fixation with 4% (v/v) paraformaldehyde at 25°C for 15 min, the cells were incubated with 300 nM of DAPI at 25°C for 30 min.

### Image acquisition

The sample containing cells was placed on the stage of a Nikon ECLIPSE Ti inverted fluorescence microscope equipped with a CFI plan fluor 10×/0.30 N.A. objective lens (Nikon, Tokyo, JAPAN), CCD camera (ORCA-R2; Hamamatsu Photonics, Hamamatsu City, JAPAN), mercury lamp (Intensilight; Nikon), XYZ automated stage (Ti-S-ER motorized stage with encoders; Nikon), and filter cubes for fluorescence channels (DAPI and GFP HYQ; Nikon).

## Results

### COP-VUV-MPS showed resistance to acid, base and organic solvent treatments

To evaluate the resistance of COP by organic solvents, we treated COP with several solvents and measured the changes in the size (**Fig. 2**). COP showed only less than 0.5% of deformity after chemical treatments. Thus, COP showed its advantages over the PDMS-based MPS, especially with regard to chemical resistance to deformation against a variety of solvents.

**Fig. 2.**
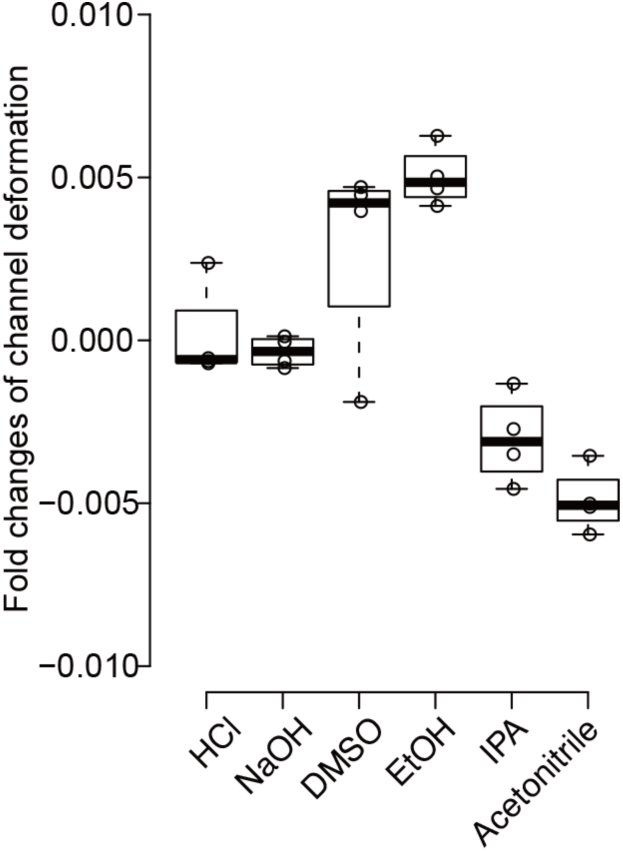
Stable structures of cyclo olefin polymer-vacuum ultraviolet-microphysiological system (COP-VUV-MPS) after chemical treatments. A COP-VUV-MPS showed chemical resistance against acid (1 M HCl), base (1 M NaCl) and organic solvents (dimethyl sulfoxide [DMSO], ethanol [EtOH], isopropanol [IPA] and acetonitrile) in terms of deformation of microstructures and assembly. Center lines show the medians; box limits indicate the 25th and 75th percentiles; whiskers extend 1.5 times the interquartile range from the 25th and 75th percentiles, outliers are represented by dots; data points are plotted as open circles. n = 4 sample points.

### COP-VUV-MPS did not absorb hydrophobic molecules

The molecule absorption into PDMS caused issues not only for cell-based assays but also for drug development. Therefore, we evaluated the absorption of molecules by the tested materials. As shown in **Fig. 3a and b**, respectively, although COP and VUV-treated COP showed slightly higher fluorescent signals in both AdipoRed and DXR than glass, they showed dramatically lower fluorescent signals than PDMS. Thus, these results suggested that COP materials showed a significant reduction of absorption of the tested compounds and can be utilized for future cell culture experiments and drug discovery.

**Fig. 3.**
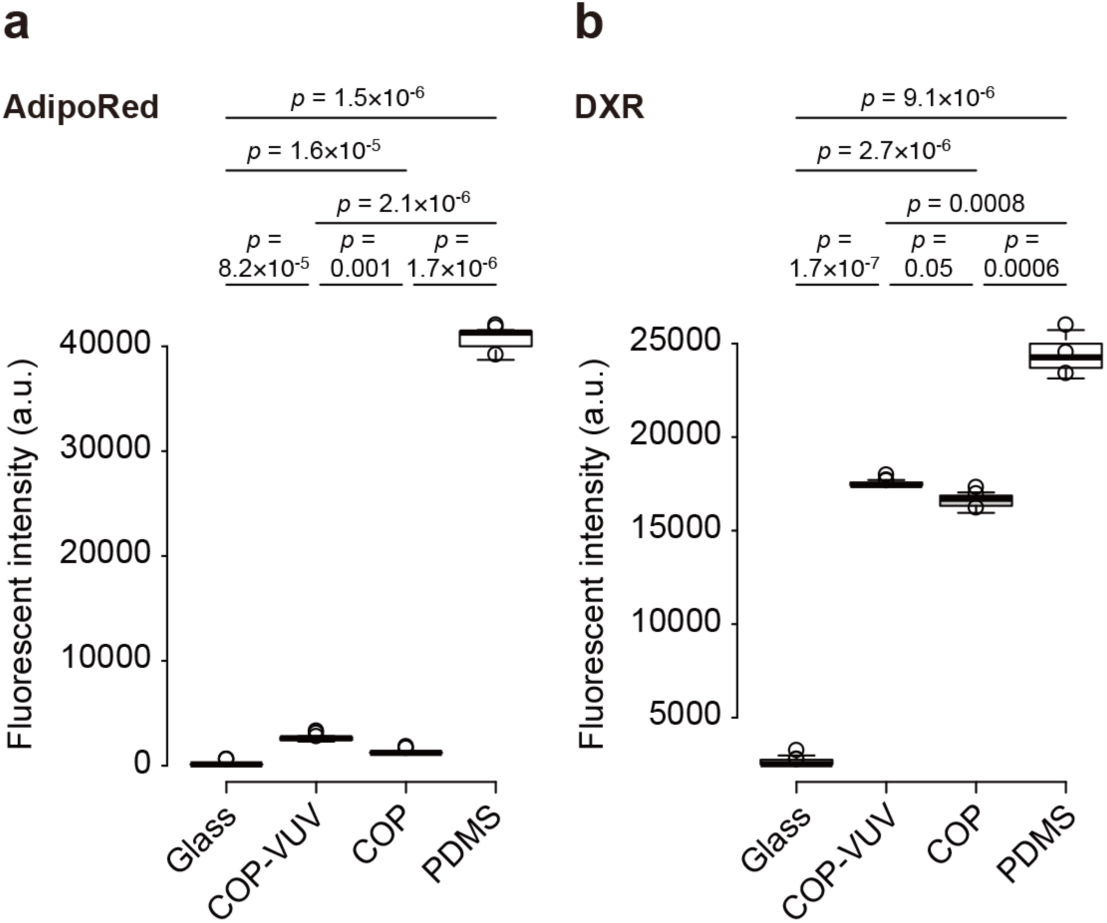
Evaluation of molecule absorption in materials used in microphysiological systems (MPSs). (a, b) Cyclo olefin polymer (COP) shows less absorption of hydrophobic compounds {a, AdipoRed (ex. 553 nm; em. 637 nm) and b, doxorubicin (ex. 470 nm; em. 585 nm)} than that of PDMS. Glass shows no absorption of these compounds. Center lines show the medians; box limits indicate the 25th and 75th percentiles; whiskers extend 1.5 times the interquartile range from the 25th and 75th percentiles, outliers are represented by dots; data points are plotted as open circles. n = 3 sample points.

### COP-VUV-MPS allowed cultivation and self-renewal of hPSCs without undesired differentiation, compared to other MPSs

To demonstrate the feasibility of a COP-VUV-MPS for cell culture and analyses, we demonstrated the culture of undifferentiated hPSCs, which have high sensitivity to the changes in the cell culture environment, and examined how such changes altered their phenotypes and gene expression (**Fig. 4**). As a comparison with COP-VUV-MPS, we used COP with solvent bonding (COP-S), PDMS-MPS and, 96-well plates (Well). First, to evaluate the pluripotent status marker in 253G1 hiPSCs cultured for 5 days, immunocytochemistry was performed to measure the expression of OCT4 (or POU5F1) (**Fig. 4a**). 253G1 hiPSCs cultured in COP-VUV-MPS and PDMS-MPS did not show a noticeable difference in OCT4 expression. Then, gene expression analysis for pluripotency-associated genes (*LIN28A, NANOG, POU5F1* and *TERT*) and differentiation for the three germ layers (e.g., *PAX6*: ectoderm, *SOX17*: endoderm and *T*: mesoderm) were performed (**Fig. 4b**). COP-S-MPS showed a relatively low expression of *NANOG* and *POU5F1* compared with the others. Furthermore, the expression of *SOX17* and *T* in PDMS-MPS was significantly higher than those of the other platforms. These results suggested that COP-VUV-MPS could maintain stem cell status during the culture, compared with other platforms.

**Fig. 4.**
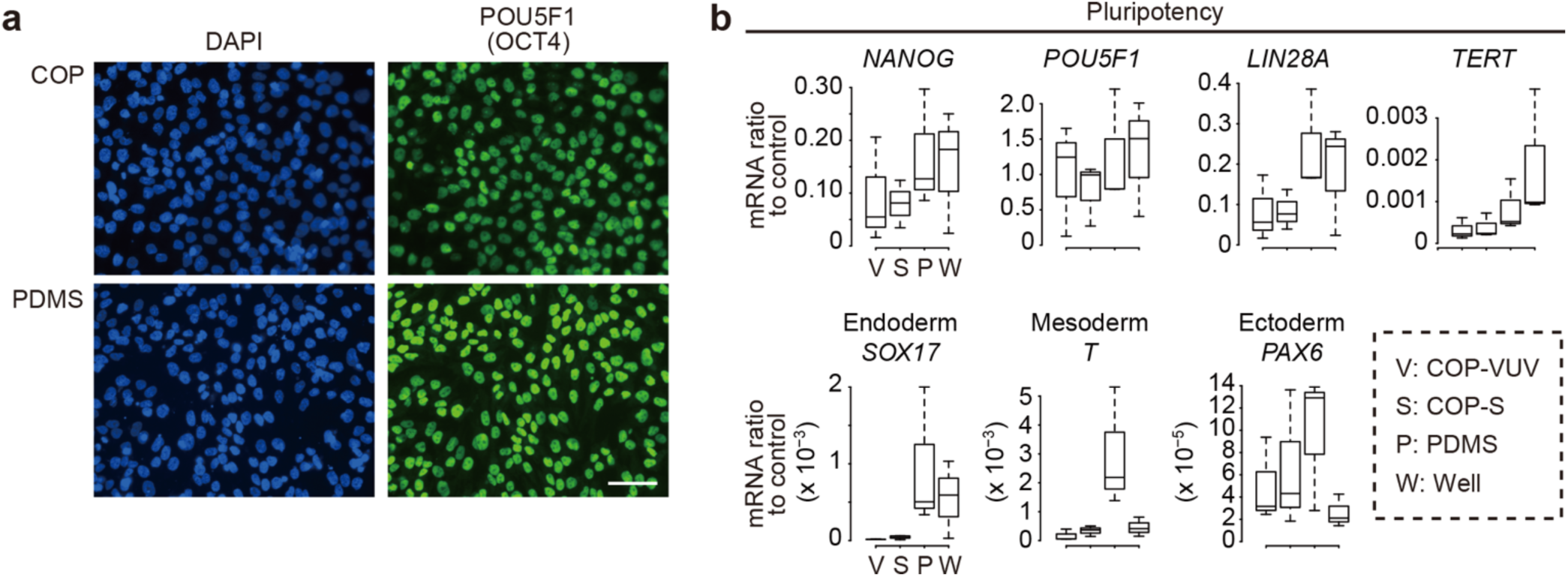
A cyclo olefin polymer-vacuum ultraviolet (COP-VUV) device allows the elimination of additional materials, which might cause cell damage. (a) Fluorescent micrographs of 253G1 human pluripotent stem cells (hPSCs) cultured in COP-VUV-microphysiological system (MPS) and polydimethylsiloxane (PDMS)-MPS, immunostained with OCT4 and DAPI. Scale bar represents 50 μm. (b) Quantitative RT-PCR to evaluate the gene expression levels associated with pluripotency (*LIN28A, NANOG, POU5F1* and *TERT*) and differentiation for three germ layers (e.g., *PAX6*: ectoderm, *SOX17*: endoderm and *T*: mesoderm). The gene expression values of GAPDH were used as an internal control. Center lines of boxplots show the medians; box limits indicate the 25th and 75th percentiles; whiskers extend 1.5 times the interquartile range from the 25th and 75th percentiles.

### COP-VUV-MPS prevented apoptosis of self-renewing hPSCs

Cell damage was evaluated via observing Annexin V staining, the marker of apoptotic cells (**Fig. 5a**). 253G1 hiPSCs cultured in a COP-S-MPS showed significantly higher apoptotic cells and stronger fluorescent intensities than those cultured in a COP-VUV-MPS. Furthermore, to investigate the apoptotic pathways altered by the tested MPS, gene expression analysis by quantitative RT-PCR was carried out for Bcl-2-associated death promoter (*BAD*), tumor protein p53 (*TP53*), Caspase 8 (*CAS8*) and *FAS* (**Fig. 5b**). These results revealed that *CAS8* expression of the cells in COP-S was dramatically increased. The two devices differed only in the bonding methods, and these results suggested that leakage of solvent from the COP-S-MPS caused apoptosis during culture.

**Fig. 5.**
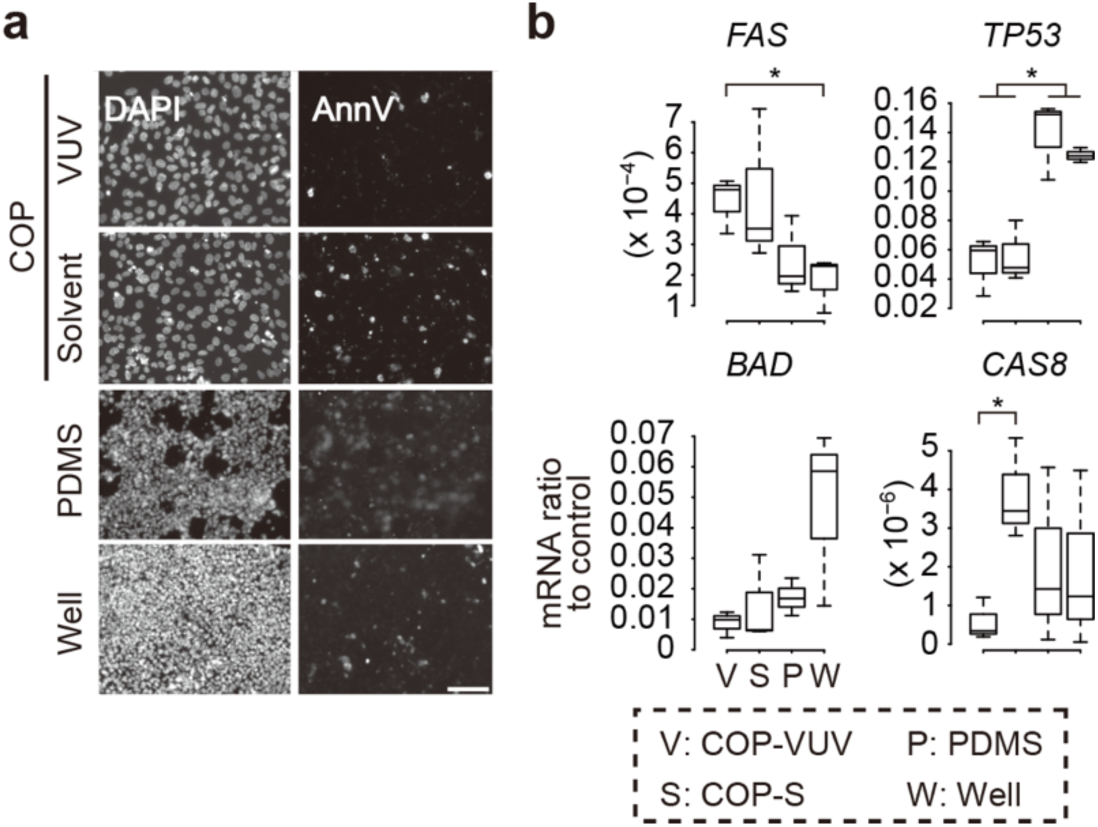
A cyclo olefin polymer-vacuum ultraviolet-microphysiological system (COP-VUV-MPS) prevents apoptosis of undifferentiated human-induced pluripotent stem cells (hiPSCs). (a) Phase contrast and fluorescent micrographs of 253G1 hPSCs cultured in COP-VUV-, COP-S- and polydimethylsiloxane (PDMS)-MPS, as well as a 96-well plate, stained with Annexin V-Alexa568 apoptosis marker (AnnV) and DAPI. Scale bar represents 20 μm. (b) Quantitative RT-PCR to evaluate the gene expression levels associated with apoptosis signaling pathways (*BAD, TP53, CAS8* and *FAS*). Center lines of boxplots show the medians; box limits indicate the 25th and 75th percentiles; whiskers extend 1.5 times the interquartile range from the 25th and 75th percentiles.

## Discussion

MPS platforms are promising for drug development, toxicological testing to predicting human responses against the drug candidates, and as alternatives to animal testing. However, several problems have to be resolved before their realization, such as MPS materials, fabrication processes less harmful for cells and mass production, and cell sources. In order to solve these problems, it is necessary to consider them comprehensively, not individually. Here, we show a solution for the advancement of the MPS platform with a low-cost plastic material that is capable of mass production and has fewer interferences for cultured cells.

PDMS has been commonly used as a material for some microfluidic devices and MPS platforms until now. Photoresists (i.e., SU-8) are also used for not only molds but also microfluidic structures. Although these materials are very convenient to fabricate at the lab with soft-lithography, ingredients in these materials may leak from the devices, affecting cellular responses and assays. Indeed, we have previously reported that the microfabrication materials used in microfluidic cell culture devices cause gene expression changes, affecting the differentiation process.^25^ Moreover, PDMS and photoresists may not be suitable for the mass production of MPS. Thus, more stable materials are required to improve MPS platforms. Glass can also be used for cell culture. However, glass requires a relatively reactive process for fabricating microfluidic devices and bonding reagents to assemble, is quite fragile for handling, and not suitable for mass production. Therefore, we used COP plastic materials for fabricating microfluidic devices. COP has several advantages over the aforementioned materials such as chemical/physical stability and high purity and optical clarity for cell culture and assays. Since COP also shows thermostability with high modulus, the metal molding process can be used for mass production with high reproducibility.

In addition to MPS materials, we also need to consider the effects of reagents/materials used for assembling MPSs. The assembly processes for glass, PS and COP often require glues or double-side tapes. Although COP shows promising properties for cell culture and assays, it also requires glues or double-side tape. Otherwise, it releases reagents into the cell culture medium, thereby influencing cellular response and assays. Thus, it is necessary to directly assemble the microfluidic structures to eliminate the concerns regarding leakage of undesired materials. Our proposed method, photobonding, with the use of VUV excimer lamp, allowed direct bonding of COP materials. The COP microfluidic device with photobonding showed high mechanical stability, and the COP material had to be destroyed to dissemble the microfluidic device. Thus, we confirmed that COP-VUV-MPS allowed the culture and self-renewal of hPSCs without undesired cellular damage.

## Conclusions

We have developed a mass-productive MPS with the use of COP materials and VUV photobonding process to eliminate the previous concerns of commonly used PDMS materials for MPS. This COP-VUV-MPS shows better resistance to chemical treatments and avoids molecule absorption. It also allows culturing very sensitive hiPSCs without altering undesired cellular phenotypes, as well as not altering gene expression during the culture. These characteristics are very important for the practical applications of MPS for drug discovery, and our MPS and approach will potentially fulfill the requirements of such applications.

## Acknowledgments

Funding was generously provided by the Japan Society for the Promotion of Science (JSPS; 16K14660, and 17H02083). The WPI-iCeMS is supported by the World Premier International Research Center Initiative (WPI), MEXT, Japan.

## References

1 J. Ribas, J. Pawlikowska and J. Rouwkema, Microphysiological Syst., 2018, 2, 10.

2 L. Ewart, E.-M. Dehne, K. Fabre, S. Gibbs, J. Hickman, E. Hornberg, M. Ingelman-Sundberg, K.-J. Jang, D. R. Jones, V. M. Lauschke, U. Marx, J. T. Mettetal, A. Pointon, D. Williams, W.-H. Zimmermann and P. Newham, Annu. Rev. Pharmacol. Toxicol., 2018, 58, 65–82.

3 U. Marx, ALTEX, 2020, 37, 1–30.

4 K. Ronaldson-Bouchard and G. Vunjak-Novakovic, Cell Stem Cell, 2018, 22, 310–324.

5 A. Guan, P. Hamilton, Y. Wang, M. Gorbet, Z. Li and K. S. Phillips, Nat. Biomed. Eng., 2017, 1, 1–10.

6 R. Abdalkader and K. Kamei, Lab Chip, 2020, 20, 1410–1417.

7 K. Kamei, Y. Kato, Y. Hirai, S. Ito, J. Satoh, A. Oka, T. Tsuchiya, Y. Chen and O. Tabata, RSC Adv., 2017, 7, 36777–36786.

8 V. van Duinen, S. J. Trietsch, J. Joore, P. Vulto and T. Hankemeier, Curr. Opin. Biotechnol., 2015, 35, 118–126.

9 M. Mehling and S. Tay, Curr. Opin. Biotechnol., 2014, 25, 95–102.

10 B. J. van Meer, H. de Vries, K. S. A. Firth, J. van Weerd, L. G. J. Tertoolen, H. B. J. Karperien, P. Jonkheijm, C. Denning, A. P. IJzerman and C. L. Mummery, Biochem. Biophys. Res. Commun., 2017, 482, 323–328.

11 M. W. Toepke and D. J. Beebe, Lab Chip, 2006, 6, 1484.

12 E. Berthier, E. W. K. Young and D. Beebe, Lab Chip, 2012, 12, 1224.

13 C.-W. Tsao, Micromachines, 2016, 7, 225.

14 S. Puza, E. Gencturk, I. E. Odabasi, E. Iseri, S. Mutlu and K. O. Ulgen, Biomed. Microdevices, 2017, 19, 40.

15 L. Yi, W. Xiaodong and Y. Fan, J. Mater. Process. Technol., 2008, 208, 63–69.

16 A. M. D. Wan, T. A. Moore and E. W. K. Young, J. Vis. Exp., 2017, 119, e55175.

17 J. A. Thomson, J. Itskovitz-Eldor, S. S. Shapiro, M. A. Waknitz, J. J. Swiergiel, V. S. Marshall and J. M. Jones, Science, 1998, 282, 1145–7.

18 J. Yu, M. A. Vodyanik, K. Smuga-Otto, J. Antosiewicz-Bourget, J. L. Frane, S. Tian, J. Nie, G. A. Jonsdottir, V. Ruotti, R. Stewart, I. I. Slukvin and J. A. Thomson, Science, 2007, 318, 1917–1920.

19 K. Takahashi, K. Tanabe, M. Ohnuki, M. Narita, T. Ichisaka, K. Tomoda and S. Yamanaka, Cell, 2007, 131, 861–872.

20 D. Park, J. Lim, J. Y. Park and S.-H. Lee, Stem Cells Transl. Med., 2015, 4, 1352–1368.

21 U. Kogelschatz, Pure Appl. Chem., 1990, 62, 1667–1674.

22 G. Chen, D. R. Gulbranson, Z. Hou, J. M. Bolin, V. Ruotti, M. D. Probasco, K. Smuga-Otto, S. E. Howden, N. R. Diol, N. E. Propson, R. Wagner, G. O. Lee, J. Antosiewicz-Bourget, J. M. C. Teng and J. A. Thomson, Nat. Methods, 2011, 8, 424–429.

23 K. Watanabe, M. Ueno, D. Kamiya, A. Nishiyama, M. Matsumura, T. Wataya, J. B. Takahashi, S. Nishikawa, S. Nishikawa, K. Muguruma and Y. Sasai, Nat. Biotechnol., 2007, 25, 681–686.

24 K. Kamei, M. Ohashi, E. Gschweng, Q. Ho, J. Suh, J. Tang, Z. T. For Yu, A. T. Clark, A. D. Pyle, M. A. Teitell, K.-B. Lee, O. N. Witte and H.-R. Tseng, Lab Chip, 2010, 10, 1113–1119.

25 K. Kamei, Y. Hirai, M. Yoshioka, Y. Makino, Q. Yuan, M. Nakajima, Y. Chen and O. Tabata, Adv. Healthc. Mater., 2013, 2, 287–291.

